# Decreased torsinA or LINC Complex Function Rescues a Laminopathy in *Caenorhabditis elegans*

**DOI:** 10.1101/2020.04.22.055988

**Authors:** Gabriela Huelgas-Morales, Mark Sanders, Gemechu Mekonnen, Tatsuya Tsukamoto, David Greenstein

## Abstract

The function of the nucleus depends on the integrity of the nuclear lamina, an intermediate filament network associated with the linker of nucleoskeleton and cytoskeleton (LINC) complex spanning the nuclear envelope. In turn, the AAA+ ATPase torsinA regulates force transmission from the cytoskeleton to the nucleus. In humans, mutations affecting nuclear envelope-associated proteins cause laminopathies, including progeria, myopathy, and dystonia. We report that decreasing the function of the *C. elegans* torsinA homolog, OOC-5, rescues the sterility and premature aging caused by a null mutation in the single worm lamin homolog, *lmn-1*. Loss of OOC-5 activity prevents nuclear collapse in *lmn-1* mutants by disrupting the function of the LINC complex. These results suggest that LINC complex-transmitted forces damage nuclei with a compromised nuclear lamina.

**One Sentence Summary:** Inhibiting LINC complex activity prevents a progeric syndrome in *C. elegans*.

## Main Text

Deleterious mutations in many conserved nuclear envelope (NE) components cause human diseases (*1*). Despite the widespread expression of NE components, these diseases often cause tissue-specific pathologies. For example, mutations affecting the NE-associated AAA+ ATPase torsinA, which is expressed in most cell types, cause early-onset DYT1 dystonia and congenital joint contractures (*2-4*). In the nematode *C. elegans*, mutations in the torsinA ortholog, OOC-5, cause oocyte growth and embryonic polarity defects (*5, 6*). The cellular functions of torsinA appear to be conserved across cell types and species because distantly related organisms lacking torsinA function display similar ultrastructural NE defects (*7, 8*). Nonetheless, the molecular interactions by which torsinA carries out its cellular functions remain enigmatic.

To address the mechanism of OOC-5 function, we purified OOC-5-associated proteins from soluble and membrane protein fractions using tandem affinity purification (fig. S1 and table S1). Both purifications identified highly conserved regulators of endoplasmic reticulum (ER) and NE homeostasis (table S1). The OOC-5-associated proteins include OOC-3, a protein required for OOC-5 localization and function (*5, 6*), ER chaperone CNX-1/calnexin, which associates with torsinA in mammalian cells (*9*), and GEI-18 and T09B4.2, the divergent *C. elegans* homologs of the torsinA-interacting proteins LAP1 and LULL1 (*10*). Also among the OOC-5-associated proteins was SUN-1, the SUN domain inner nuclear membrane protein component of LINC complexes in the *C. elegans* germline (*11*). SUN domain proteins interact with nuclear lamins (*12, 13*), and the sole *C. elegans* lamin, LMN-1 (*14*), was well represented among OOC-5-associated proteins (42-43% coverage). The genetic interaction studies described below suggest that the association between OOC-5, which resides in the lumen of the NE and ER, and nucleoplasmic LMN-1 is likely mediated through SUN-1.

To address the significance of the protein associations with OOC-5, we conducted genetic analyses. The most striking result was an interaction between *ooc-5* and *lmn-1*. Most *lmn-1* null mutants are agametic, sterile, and have small gonads (Fig. 1B and Fig. 2B). However, 12.3 ± 7.8% of animals (“escapers”) develop larger gonads and produce dead embryos (Fig. 1C and Fig. 2B). Surprisingly, compromising *ooc-5* function rescued the sterility of *lmn-1* mutants such that all animals produced gametes and dead embryos (Fig. 1D, Fig. 2, C and D, and fig. S2). Importantly, reducing the dosage of *ooc-5* was sufficient to suppress *lmn-1* sterility. Reducing or eliminating *ooc-5* function also suppressed the progeric defects (Fig. 1E) previously described in *lmn-1(–)* mutants (*15*).

**Fig. 1.**
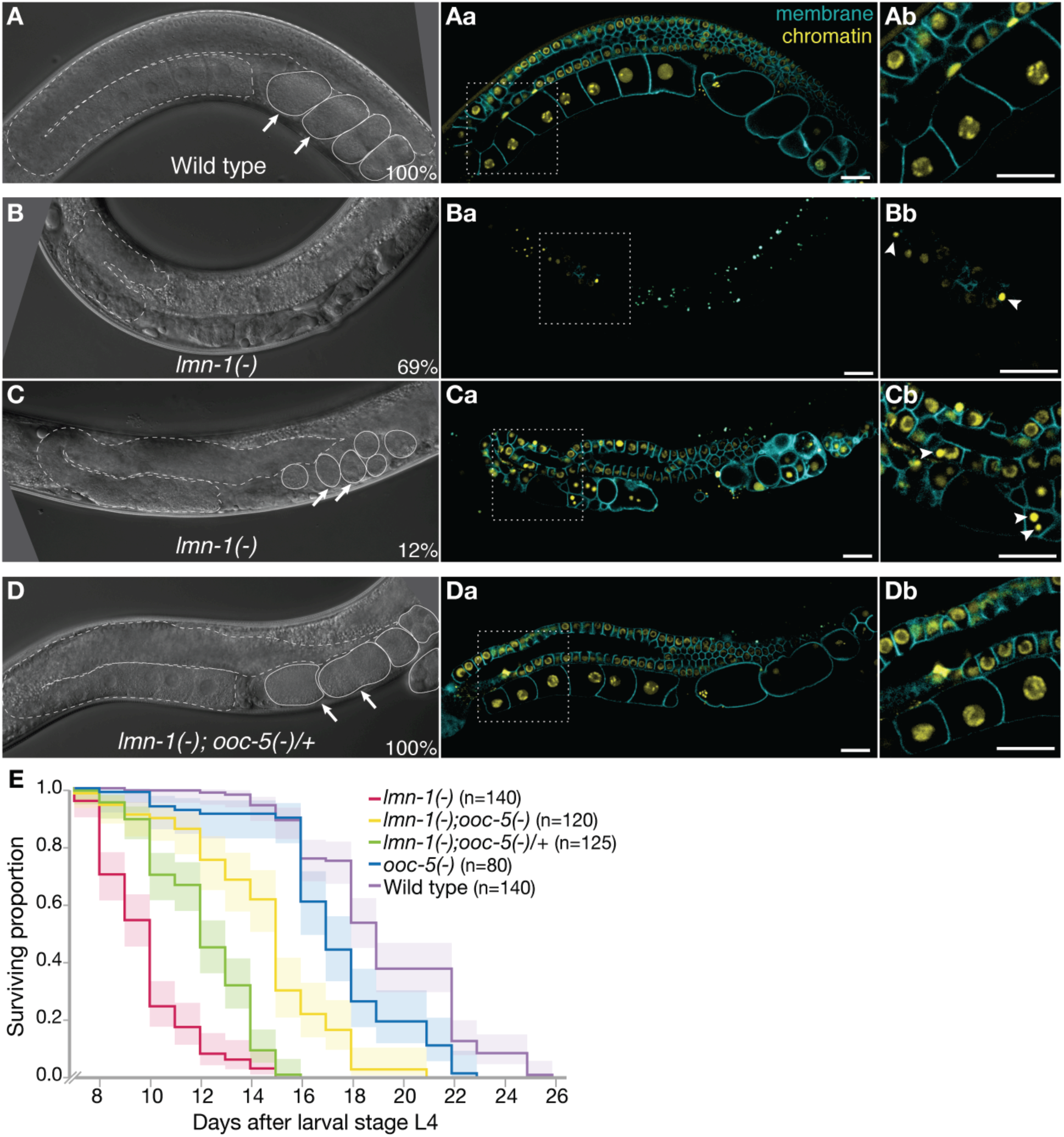
Loss of *ooc-5* rescues defects in lamin mutants. (**A**-**D**) DIC images of adult hermaphrodites. Arrows indicate embryos. (**Aa**-**Da**) Fluorescence micrographs of adult hermaphrodites. Cellular membranes of germ cells are visualized using a GFP::PH domain fusion and chromatin is visualized with a histone H2B::mCherry fusion. (**Ab**-**Db**) Insets are twice the size of the dotted-line region. Arrowheads indicate collapsing nuclei. Bar, 20 µm. (**E**) Kaplan-Meier survival curves of the indicated genotypes. Three replicates were conducted, and confidence intervals are shadowed.

**Fig. 2.**
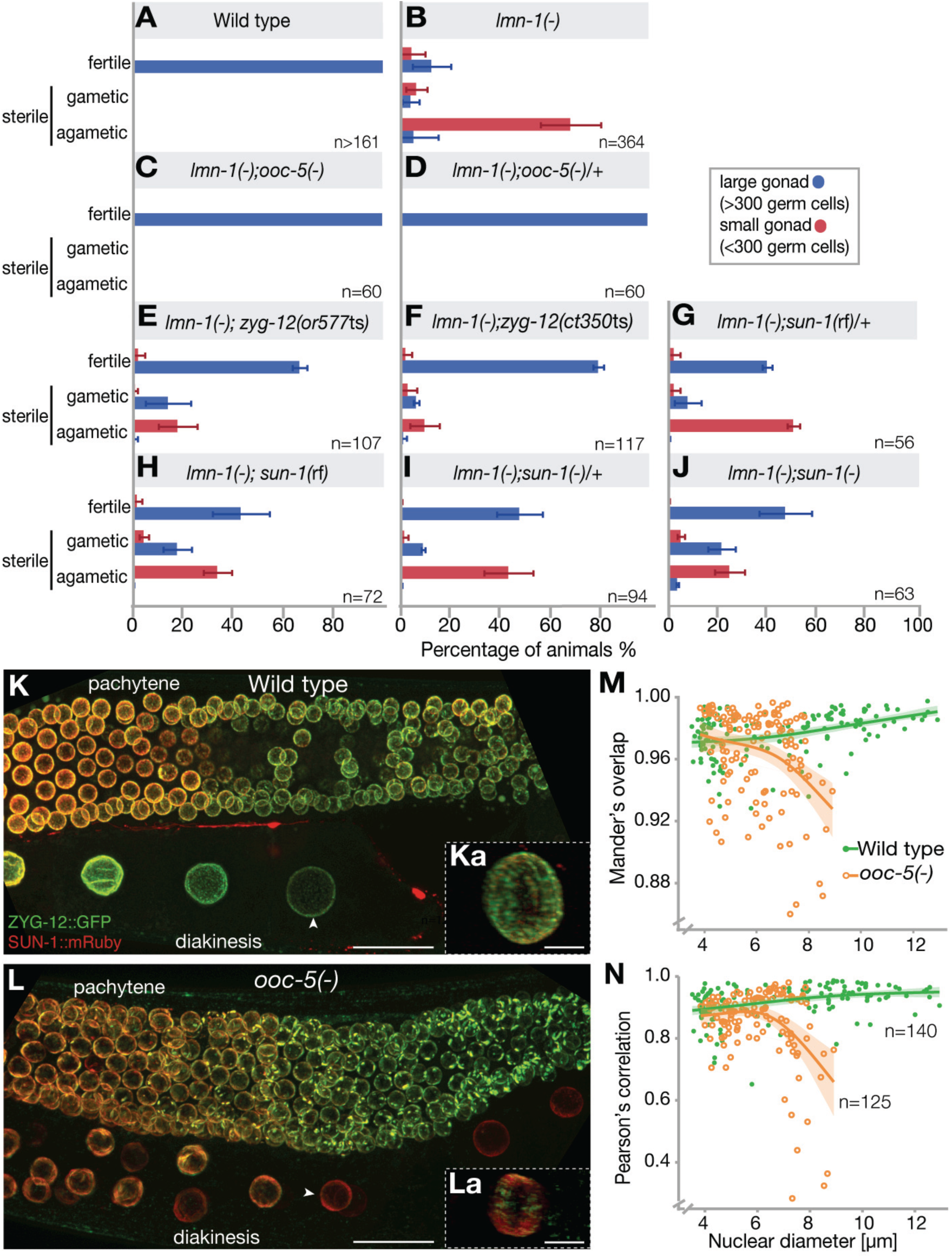
*ooc-5* regulates the LINC complex. (**A**-**J**) Hermaphrodites of the indicated genotypes grown at 20°C, were classified as fertile (≥1 embryo), sterile gametic (sperm/oocytes), or sterile agametic. Animals with more than 300 germ cells in a gonad arm were further classified as having large (blue) versus small gonads (red). (**K** and **L**) Maximum projections of Z-stack super-resolution confocal images of ZYG-12::GFP and SUN-1::mRuby in the gonad. Bar, 20 µm. (**Ka** and **La**) 3D-reconstructions of the nuclei indicated with arrowheads. Bar, 5 µm. (**M** and **N**) Colocalization of ZYG-12::GFP and SUN-1::mRuby in the NE. Bootstrap confidence regions are shadowed.

The germ cells of the *lmn-1* escapers appeared to deteriorate as they progressed through meiotic prophase; they assembled a synaptonemal complex (SC) but their nuclei collapse soon after the pachytene stage (fig. S3). SUN-1 lost its NE localization and appeared diffuse in most collapsing nuclei (fig. S4). Diakinesis-stage chromosomes were rare in *lmn-1* mutant oocytes but were observed in most or all *lmn-1* mutants after reducing *ooc-5* function (Fig. 1 and fig. S3). Decreasing *ooc-5* function did not appear to rescue the elevated levels of DNA damage observed in the late-stage pachytene female germ cells of *lmn-1* mutants (fig. S5).

Studies in mammalian cells show that torsinA regulates force transmission from the cytoplasm to the nucleus (*16*). Thus we examined a model in which OOC-5 promotes LINC complex function, which damages nuclei with a compromised nuclear lamina. We tested for genetic interactions between *lmn-1(–)* mutants and genes encoding components of the LINC complex, utilizing mutations in *sun-1* and *zyg-12*, which encodes the KASH domain outer nuclear membrane LINC complex-component in the germline (*11*). As observed for *ooc-5*, the proportion of fertile *lmn-1* animals increased when the LINC complex function was compromised, and nuclear collapse was substantially ameliorated (Fig. 2, E–J, fig. S2, and fig. S6). These results suggest that OOC-5 and the LINC complex-transmitted forces contribute to nuclear damage in *lmn-1* mutants.

To assess whether OOC-5 promotes LINC complex assembly, we used super-resolution microscopy to examine the colocalization of SUN-1 and ZYG-12 in the wild type and *ooc-5* mutants. Prior work showed discontinuities in ZYG-12 localization in *ooc-5* mutants commencing in pachytene germ cells (*8*). In addition to these discontinuities, we observed a reduction of ZYG-12 NE localization as germ cells progressed from the pachytene stage to the diakinesis stage of meiotic prophase (Fig. 2L). By contrast, SUN-1 remained in the NE at high levels in *ooc-5* mutants (Fig. 2L). We quantified SUN-1 and ZYG-12 colocalization as a function of nuclear diameter and observed that as the germ cells grew, colocalization declined (Fig. 2, M and N). These results suggest that OOC-5 promotes the assembly or maintenance of the LINC complex, which appears particularly important for growing meiotic prophase nuclei.

The LINC complex is essential for the pairing of homologous chromosomes in *C. elegans* meiosis (*17-19*). OOC-5 is not required for SC formation (fig. S3) and appears dispensable for meiotic chromosome segregation (*5*). To examine whether OOC-5 promotes the efficiency of meiotic pairing, we examined the localization of ZIM-3 and HIM-8, which bind the pairing center regions on chromosomes I and IV and the X chromosome, respectively (*20*). The spatial distribution of unpaired meiotic chromosomes across the distal gonad suggests that pairing of chromosomes I and IV is delayed in *ooc-5* mutants (Fig. 3B). By contrast, *ooc-5* appears dispensable for the timely pairing of the X chromosome (Fig. 3C). Persistent SUN-1 phosphorylation correlates with synaptic or recombination defects (*21*). We observed that phosphorylated SUN-1 (S8Pi) persisted in an extended zone in *ooc-5* mutants (Fig. 3, D–F), and the number of SUN-1 foci and their speed of motion increased (Fig. 3, G and H, and mov. S1–3). The increased movement of SUN-1 foci in *ooc-5* mutants may reflect partial release from cytoskeletal tension. These results suggest that the function of the LINC complex is perturbed in *ooc-5* mutants even in early prophase germ cells, where SUN-1 and ZYG-12 colocalize normally.

**Fig. 3.**
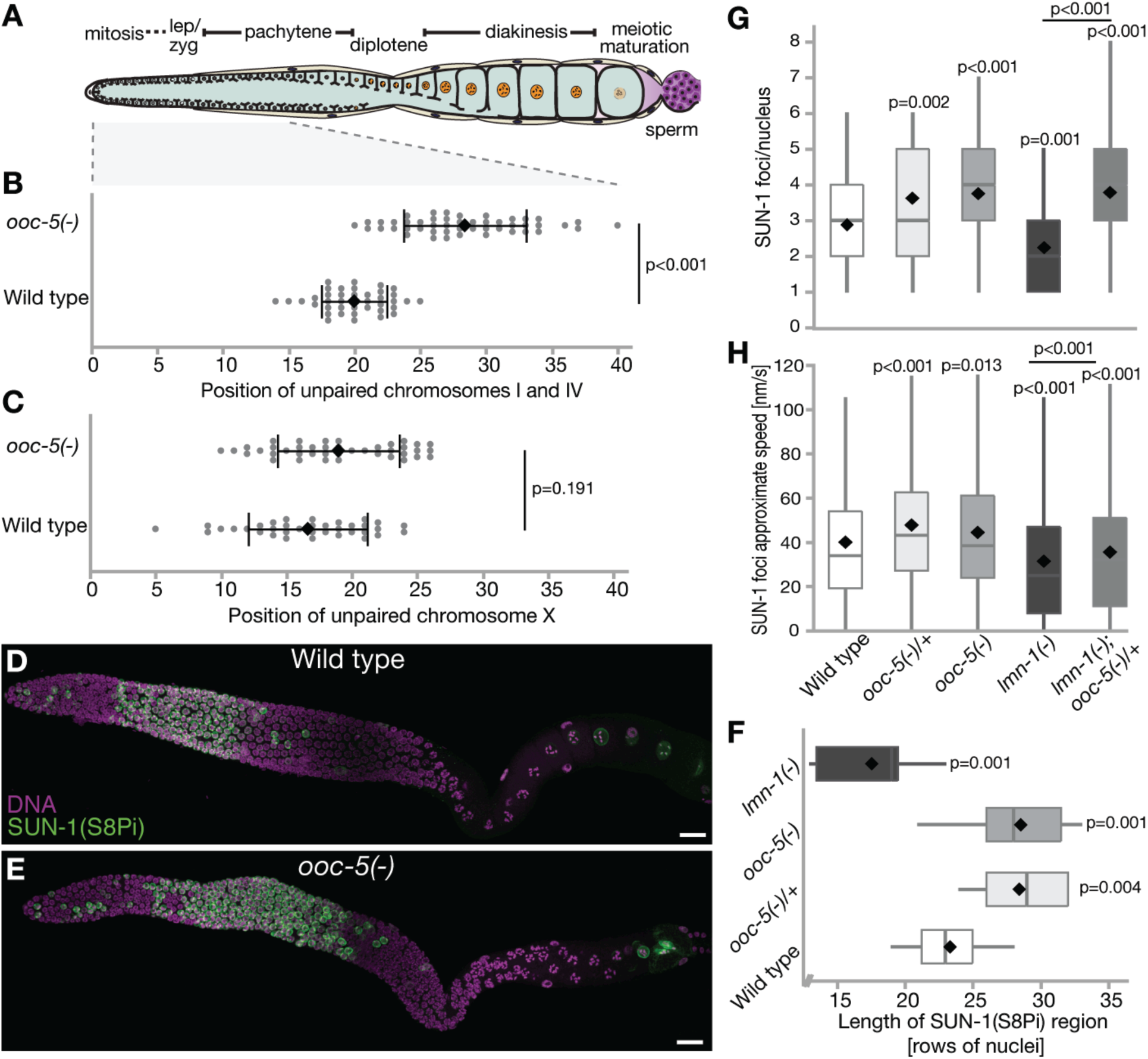
LINC complex function during meiosis is compromised in *ooc-5* mutants. (**A**) Diagram of the *C. elegans* adult hermaphrodite gonad. (**B** and **C**) Spatial distribution of unpaired chromosomes I and IV (**B**) and X chromosomes (**C**), visualized by immunostaining for ZIM-3 and HIM-8, respectively (see fig. S7). (**D** and **E**) Gonads immunostained for phosphorylated SUN-1 (S8Pi, green) and DNA (magenta). Bar, 20 µm. (**F**) Extent of the meiotic region containing phosphorylated SUN-1 (S8Pi). Diamonds indicate mean values. (**G**) Quantification of SUN-1 foci per nucleus, and (**H**) of their approximate speed of movement from time-lapse images (see mov. S1-3).

*C. elegans* OOC-5 and mammalian torsinA are required for the proper localization of nuclear pore complex (NPC) proteins (*8, 22*). Mislocalization of nucleoporins (Nups) may be sufficient to explain the embryonic polarity defects in *ooc-5* mutants (*23*). The two transmembrane Nups, NPP-12 and NPP-22, were identified among OOC-5-associated proteins (table S1). A prior study concluded that an NPP-22::GFP-expressing transgene localized more normally in *ooc-5* mutants than other Nups, but the localization of NPP-12 was not examined (*8*). Thus, we used genome editing to generate GFP fusions to the endogenous proteins and analyzed their localization. Both NPP-12::GFP and GFP::NPP-22 exhibited substantial mislocalization in *ooc-5* mutants, accumulating largely outside the NE and in the cytoplasm of proximal oocytes (Fig. 4, B, D, and G). This mislocalization of Nups in *ooc-5* mutants might be a consequence of the impaired function of the LINC complex, as prior work showed that Sun1 disruption in mammalian cells lead to NPC mislocalization (*24*). Consistent with this possibility, NPP-12::GFP exhibited mislocalization in *sun-1* and *zyg-12* mutants (Fig. 4, E–G).

**Fig. 4.**
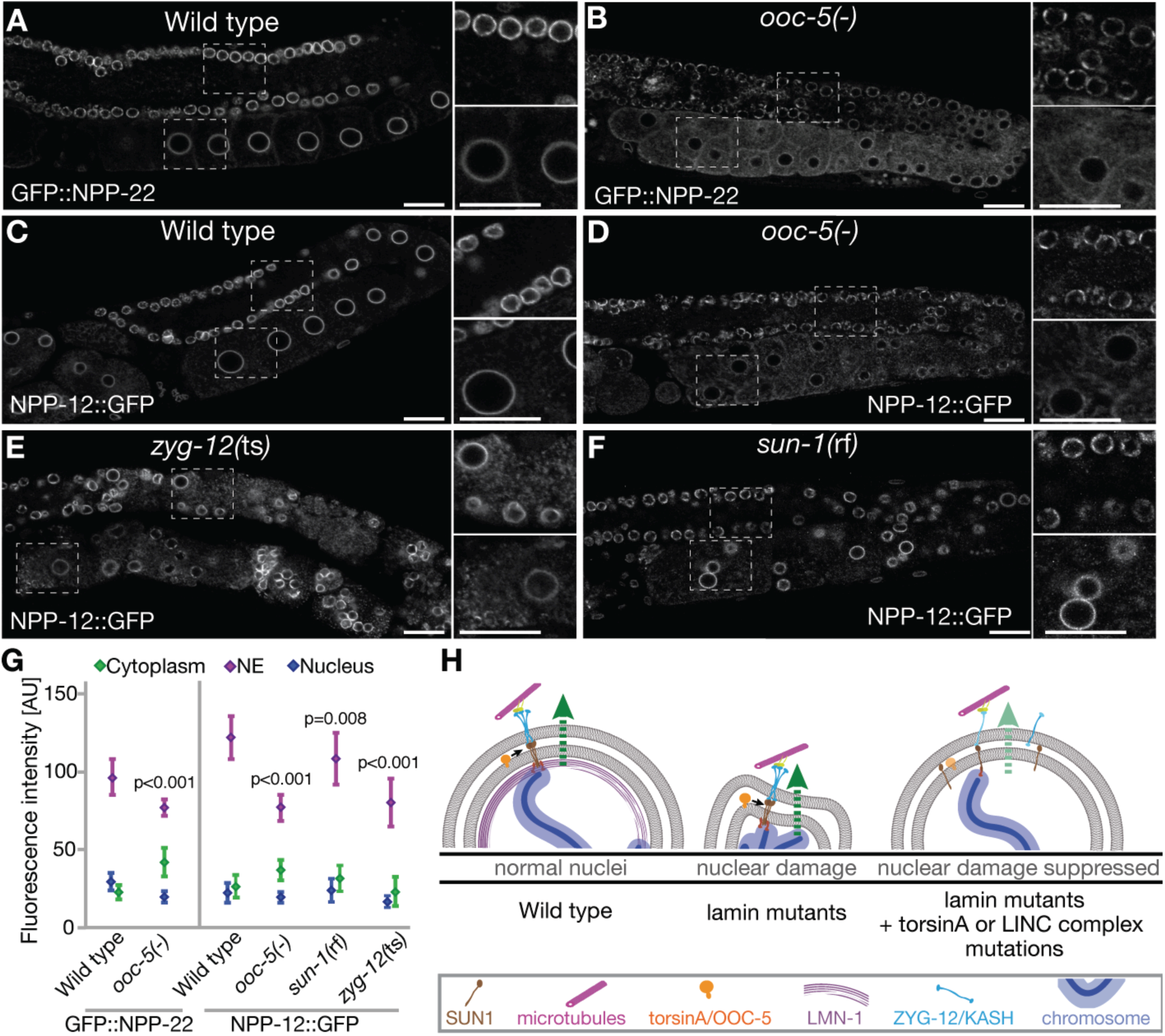
OOC-5 and the LINC complex promote the localization of transmembrane Nups. (**A**-**F**) Confocal images of the expression of GFP::NPP-22 or NPP-12::GFP in the gonad. Animals were grown at 20°C except *zyg-12*(ts) mutants, which were grown at 25°C. Bars, 20 µm. (**G**) Quantification of the fluorescence intensity of GFP::NPP-22 or NPP-12::GFP in subcellular compartments (mean ± SD). (**H**) Model. OOC-5 promotes LINC complex function, damaging *lmn-1* mutant nuclei. Green arrows indicate force.

Our results indicate that OOC-5 promotes LINC complex functions needed to maintain NE homeostasis. Further, these data suggest that in the absence of a functional nuclear lamina, LINC complex-dependent forces promoted by OOC-5 result in nuclear damage (Fig. 4H). Consistent with this model, disruptions of the LINC complex can ameliorate defects in cultured cell and mouse models of Hutchinson-Gilford progeria (*25*). Also, LINC complex-dependent forces may be important for maintaining nuclear structure and protecting the NE from mechanical damage, as suggested by a study of fibroblasts from DYT1 dystonia patients (*26*). Age-related alterations in the ability to cope with mechanotransduction across the NE may contribute to progeric syndromes and reduce normal healthspan.

## Acknowledgments

We thank G. W. Gant Luxton for encouraging us to pursue this project. We thank Abby Dernburg, Verena Jantsch, Sarit Smolikove, Daniel Starr, and Anne Villeneuve for providing antibodies. We thank Jack Powers, Caroline Spike, and Todd Starich for comments on the manuscript and encouragement. We thank Ana E. Garcia-Vedrenne for guidance on statistics. We thank Nikon Instruments for use of the SoRa microscope. Some strains were provided by the Caenorhabditis Genetics Center.

## Author contributions

G. H.-M. and D. G. conceptualized the research and the experimental strategy. G. H.-M. and T. T. defined OOC-5-associated proteins. G. H.-M. and G. M. generated the *C. elegans* strains and conducted genetic analyses. G. H.-M and M. S. conducted the microscopy and quantified the results. G. H.-M. and D. G. wrote the manuscript. All authors contributed to data interpretation and commented on the manuscript.

## Data and materials availability

All data supporting the conclusions of this study are present in the paper or the Supplemental Materials. *C. elegans* strains are available upon request.

## Supplementary Materials

### Materials and Methods

#### Strains

The genotypes of strains used are reported in the Supplement, table S4. Genes and mutations are described in WormBase (www.wormbase.org). The following mutations were used: LGI-*lmn-1(tm1502), npp-12(tn1780 [npp-12::gfp::tev::3xflag]), T09B4.2(tn1886)*; LGII-*ooc-5(tn1757), ooc-5(it145) unc-4(e120), ooc-5(tn1709), ooc-5(tn1758[ooc-5::gfp::tev::3xflag]), zyg-12(or577*ts*), zyg-12(ct350*ts); LGIV–*gei-18(tn1874)*; LGV–*sun-1(jf18), sun-1(gk199), npp-22(tn1794[gfp::tev::3xflag::npp-22])*. The following rearrangements were used: *tmC18[dpy-5(tmIs1236)]* I, *mIn1[dpy-10(e128)mIs14]* II, and *tmC12[egl-9(tmIs1194)]* V. The following transgenes and transgene insertions were used: *ieSi21[sun-1p::sun-1::mruby::sun-13’UTR+Cbr unc-119, itIs37[pie-1p::mCherry::H2B::pie-1 3’UTR+unc-119(+)]* IV, *ojIs9[zyg-12(all)::gfp+unc-119(+)]*, and *itIs38[pie-1p::GFP::PH(PLC1delta1)+unc-119(+)].*

#### Genome editing

CRISPR-Cas9 genome editing used pRB1017 to express single guide RNA (sgRNA) under control of the *C. elegans* U6 promoter (*27*). To generate the sgRNA plasmids, we annealed oligonucleotides and ligated them to the pRB1017 vector, previously digested with *Bsa*I. pDD162 was the source of Cas9, expressed under control of the *eft*-3 promoter (*28*).

We generated deletions in *ooc-5* using a modification of the method of Paix *et al*. 2014 (*29*). Adult hermaphrodites of the N2 strain were injected with a mix containing the sgRNA plasmids at 25 µg/ml each, pDD162 at 50 µg/ml, pmyo-2::tdTomato at 4 µg/ml, pCFJ104 at 25 µg/ml, and the oligonucleotide used as repair template at 500 nM. The oligonucleotides used as repair templates are listed in table S5. Transgenic animals from the F1 generation were singled out and pools of F2 were screened by the polymerase chain reaction (PCR). For the *ooc-5* mutations F2 siblings from positive plates were crossed into the *mIn1* balancer.

We generated N-terminal or C-terminal translational fluorescent protein gene fusions using the self-excising-cassette method (*30*). The oligonucleotides to generate the repair templates are listed in table S5. All new alleles were verified by Sanger sequencing, and outcrossed to wild-type males at least three times.

#### OOC-5 immunopurifications

OOC-5:: GFP::TEV::3xFLAG and interacting proteins were purified from the DG4377 strain. Purifications from the wild-type strain (N2) were run in parallel to use as control. The worms were grown at 15°C on peptone-enriched plates seeded with bacterial strain NA22. Embryos were isolated by alkaline hypochlorite treatment (20% bleach and 0.5 N NaOH), washed in M9 buffer, and allowed to hatch overnight in the absence of food at 15°C. 30,000 L1-stage animals were cultured on each 150 × 15 mm petri dish on peptone-enriched medium with NA22 as a food source until they reached adulthood (60-70 petri dishes per strain).

To prepare the lysate, young adults were washed thoroughly with M9 and then frozen in droplets with liquid nitrogen. Then, the droplets were ground into fine powder, which was resuspended in an equal volume of lysis buffer, which was 75 mM HEPES (pH 7.5), 1.5 mM MgCl_2_, 150 mM KCl, 15% glycerol and 1.5x protease inhibitors (cOmplete Mini EDTA-free, Roche). The crude extracts were sonicated for a total of 3 min at 30% amplitude (Fisher Scientific, Model 500 Sonic Dismembrator) on ice. The soluble fraction was prepared by ultra-centrifugation of the crude extract at 135,000 x g for 30 min at 4°C. The membrane fraction was prepared by resuspending the remaining pellet in lysis buffer plus 1% Triton x-100 (Sigma-Aldrich), sonicating for 15 s, and ultra-centrifuging at 135,000 x g for 30 min at 4°C. The protein concentration of each fraction was determined by Bradford assay (Bio-Rad protein assay dye).

For the primary immunopurification of OOC-5::GFP::TEV:: 3xFLAG, Dynabeads Protein G (Life Technologies) were crosslinked to anti-FLAG M2 antibodies (Sigma-Aldrich, F1804), according to the manufacturer’s instructions. The soluble (500 mg protein input) and membrane (170 mg protein input) fractions were incubated with these dynabeads for 1 h at 4°C with rotational mixing. Then, the beads were washed 3x with IP wash buffer, which was 50 mM HEPES (pH 7.5), 1 mM MgCl_2_, 300 mM KCl, 10% glycerol, 0.05% NP-40, 5 mM 2-mercaptoethanol, 5 mM sodium citrate, 10 µM ZnCl_2_, and protease inhibitors (cOmplete Mini EDTA-free, Roche). Immunopurified proteins were eluted with 0.17 U/ul AcTEV protease (Invitrogen) in IP wash buffer for 16 h at 4°C with rotational mixing.

For the secondary immunopurification, the Dynabeads were crosslinked to a 1:1 mixture of anti-GFP 12A6 and anti-GFP 4C9 (Developmental Studies Hybridoma Bank, University of Iowa). The TEV-eluates were incubated with these Dynabeads for 1 h at 4°C with rotational mixing. Then, the beads were washed 6x with IP wash buffer. Immunopurified proteins were eluted with 100 mM glycine (pH 2.5) and neutralized immediately with an equal volume of 2 M Tris-Cl (pH 8.5). Small aliquots of protein fractions from every step of the purifications were separated on a 4-12% Bis-Tris NuPAGE gel, transferred to Protran nitrocellulose membrane (Whatman), and it was incubated first with rabbit anti-GFP NB600-308 antibody (1:4,000) (Novus Biologicals) for 2 h, and then with peroxidase-conjugated donkey anti-rabbit antibody (1:7,000) (Jackson ImmunoResearch) (fig. S1F). Detection was performed using SuperSignal West Femto Maximum Sensitivity Substrate (Thermo Scientific).

The glycine eluates containing immunopurified proteins were concentrated on a centrifugal evaporator (Jouan RC 10-10 Vacuum Concentrator) for 5 h, briefly separated on a 12% Bis-Tris NuPAGE gel, and stained using the Colloidal Blue Staining Kit (Invitrogen) (fig. S1G). Lanes were subdivided into six slices and mass spectrometry was performed at the Taplin Biological Mass Spectrometry Facility (Harvard Medical School) using an LTQ ion-trap mass spectrometer. To account for abundant contaminant proteins that bound non-specifically to the immunopurification matrix, in addition to our control immunopurifications from the N2 strain, we utilized data from multiple purifications previously reported (*31, 32*).

#### Lifespan analysis

All the strains were maintained at 20°C. 10 mid-L4 hermaphrodites were transferred onto NGM plates freshly seeded with OP50-1. These animals were scored for viability every 1-3 days, either by movement or tapping them with a platinum wire. The wild-type animals were transferred to NGM plates from the same batch every other day for a week to avoid overcrowding by their progeny. The animals that crawled on the plate sides were censored. Kaplan-Meier survival curves were generated using the JMP Pro 14 software (SAS Institute Inc.). Wilcoxon tests were performed to individual pairs of survival curves.

#### Immunostaining

The gonads of day 1 adult hermaphrodites were dissected in Egg Salt buffer fixed in 1% paraformaldehyde for 5 min, and post fixed in cold methanol (-20°C) for 1 min. Then, the samples were blocked with 1mg/ml BSA in PBT overnight at 4°C. Primary antibody incubation was performed in PBT+BSA overnight at 4°C, while secondary antibody incubation was performed in PBT+BSA for 1.5-2 h at room temperature.

Gonads were stained with the following antibodies provided by Abby Dernburg: goat anti-SYP-1 (1:4000), rabbit anti-ZIM-3 (1:1000), guinea pig anti-HIM-8 (1:250), guinea pig anti-HTP-3 (1:300). The guinea pig anti-SUN-1(S8Pi) (1:700) was provided by Verena Jantsch, and the rabbit anti-RAD-51 (1:30,000) was provided by Sarit Smolikov. Secondary antibodies were Cy3-conjugated goat anti-mouse (1:500, Jackson ImmunoResearch), Cy3-conjugated donkey anti-chicken IgY (1:1000, Jackson ImmunoResearch), and DyLight 488-conjugated donkey anti-guinea pig (1:800, Thermofisher Scientific). 4’,6’-diamidino-2-phenylindole (DAPI) was used to detect DNA.

#### Image acquisition and processing

The images of day 1 adult hermaphrodites, anesthetized with 0.1% levamisole on 2% agarose pads, were acquired on a Nikon Ti2 inverted microscope equipped with a Plan Apo IR 60x and an HP Apo TIRF 100xH oil immersion objective lenses, NA 1.27 and 1.49 respectively. Illumination was provided by 405, 488 and 561nm lasers to the multi-point galvano and resonant scanners. The emission filters used were 450/50, 525/50 and 595/50nm. The galvano scanner was used to acquire the images. For the images shown in the figures and movies, comparable acquisition parameters were used.

To examine the colocalization of SUN-1::mRuby and ZYG-12::GFP we used Spinning Disk Super Resolution by Optical Pixel Reassignment (SoRA). 0.30 µm step z-series images were acquired in a Nikon Ti2 inverted microscope equipped with a SR HP plan apo lambda S 100X silicon oil immersion objective lens, NA 1.35. Illumination of the double-labeled samples was provided by 488 and 561nm lasers fed through a single mode fiber to the CSU-W1 SoRA spinning disk confocal head with 50-micron pinhole. The emission filters used were 522/45 and 605/70nm, respectively. Images were acquired using a Photometrics Prime SCMOS camera with a pixel size of 0.13 µm and the SoRA (1x mag) images were reconstructed. Images were deconvolved using the Nikon Elements 3D automatic deconvolution method, which utilized between 20 and 50 iterations.

The colocalization of SUN-1::mRuby and ZYG-12::GFP was quantified on images acquired with the SoRA for several nuclei from two specimens per strain, as well as on confocal images acquired with the Nikon Ti2 for several nuclei from 5 specimens per strain. The images acquired with the Nikon Ti2 were denoised and a threshold was set to include the fluorescence of SUN-1::mRuby at the nuclear envelope. Mander’s overlap and Pearson’s coefficient were determined for each nucleus at the mid focal plane, and their nuclear diameters were measured. Imaging and processing were done in Nikon Elements versions 5.10 and 5.20.

Nikon Elements GA3 was used to quantify the intensity of the fluorescence of NPP-12::GFP and GFP::NPP-22 from confocal images of day 1 adult hermaphrodite gonads. The 488 nm channel was processed as follows: the background was corrected with Rolling Ball (14 µm) function, one pass of denoising was performed, and the regional maxima and Best FocusPlane were detected. Finally, a threshold was set for identification of the features of interest. Oocyte nuclei were filtered by circularity and fluorescence intensity values were measured (mean, min and max) at the NE, inside the NE (nucleoplasm) and at the cytoplasm (rectangular region between the edge of the NE and the cell membrane) for 10 oocytes per animal, from at least 2 animals per strain. The brightness and contrast of the images shown in Figure 4 were adjusted for presentation identically across all the strains.

#### Time lapse acquisition and processing

The movement of SUN-1 foci was documented as previously reported with the following modifications (*33*). Hermaphrodites were selected as mid-L4 larvae and incubated at 20°C. About 24 h later, young adults were anesthetized in 10 mM levamisole and mounted on 2% agarose pads. 0.25 µm step z-series were acquired with the Apo IR 60x WI, 1.27 NA objective approximately every 15 s for 6-7 min per animal. Maximum intensity projections were made with ImageJ (FIJI), the images were converted to grayscale and their Lookup Tables (LUTs) inverted. Individual SUN-1 foci were followed using StackReg (*34*) and manual tracking plug-ins. The speed was determined for 84 to 390 individual foci per animal, and the number of animals assessed was equal or greater than 3 per strain.

#### Statistics

The statistical analyses described below were performed using JMP Pro v14 software (Statistical Discovery, SAS). For the suppression of sterility phenotype data, a contingency analysis of outcome by genotype was performed (see fig. S2). Standard ChiSquare tests were conducted to compare the data from each mutant to those of their respective control (e.g. *lmn-1(–);ooc-5(–)/mIn1* was compared to *lmn-1(–);mIn1*/+). SUN-1(S8Pi), SUN-1 foci number and speed of movement, and the position of unpaired chromosomes data were tested for equality of group variance. Since they all had unequal variances, non-parametric comparisons were made for each pair using the Wilcoxon method.

The comparison of fluorescence intensity of NPP-12::GFP or GFP::NPP-22 in different subcellular compartments between genetic backgrounds was performed using Prism 8 (GraphPad software, LLC.). A 2way ANOVA showed that the effect of the different genetic backgrounds was significant, P< 0.0001. Bonferroni multiple comparisons tests were performed using the fluorescent intensity of the nucleoporins in the otherwise wild-type background as controls. Figure 4G shows the p-values for these comparisons in the NE.

---

**fig. S1.**
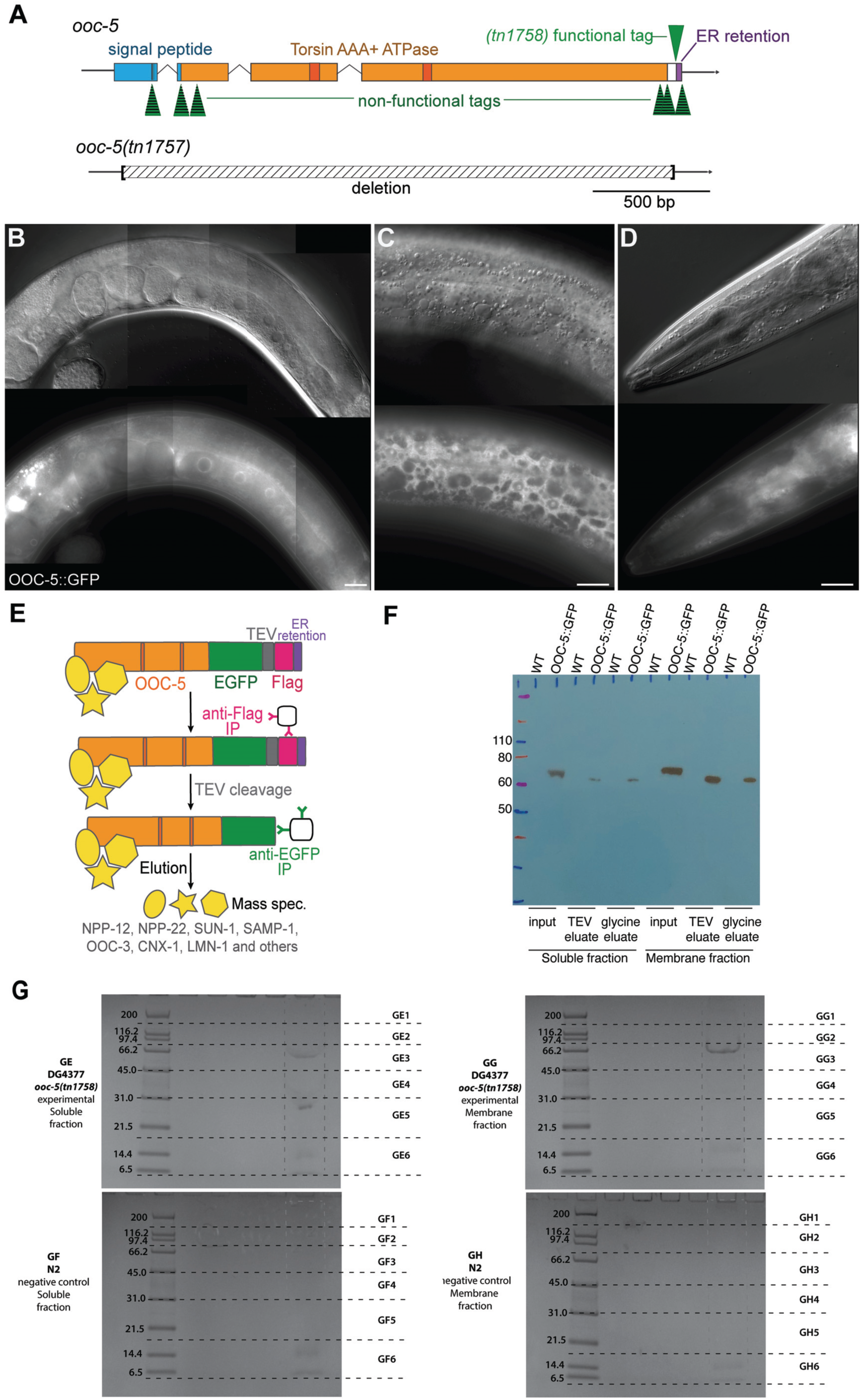
OOC-5/torsinA tandem affinity purification. (A-B) Representation of *ooc-5* locus. Above: The affinity tag was inserted in 7 different positions by genome editing, but only the insertion right before the endogenous ER retention motif (HFDDEL) exhibited functionality. The product of the new allele *ooc-5(tn1758)* is herein referred to as OOC-5::GFP. Although the OOC-5::GFP strain was viable and fertile at 15°C, it was temperature-sensitive (table S2). Thus, purifications were conducted only using animals grown at 15°C. Below: Representation of the *ooc-5(tn1757)* null allele generated by genome editing. (B-D) DIC and GFP composite images of the expression of OOC-5::GFP in the gonad (B), hypodermis (C), and head (D) of a young adult hermaphrodite grown at 15°C. Bars, 20 µm. (E) Overview of purification methods used to characterize OOC-5-associated proteins. GFP (green), tobacco etch virus (TEV) protease cleavage site (gray), and 3x Flag-tag (pink). (F) Western blot of the steps involved in OOC-5::GFP tandem affinity purification, from both the soluble and the membrane fractions. Note, the bands decreased in size slightly after TEV protease cleavage as expected, from approximately 73 to 68 KDa. (G) Coomassie staining of the OOC-5::GFP (GE and GG) and control (N2) (GF and GH) immunopurifications, for soluble and membrane fractions of the protein extracts. The lanes were cut into six slices each and processed by mass spectrometry.

**fig. S2.**
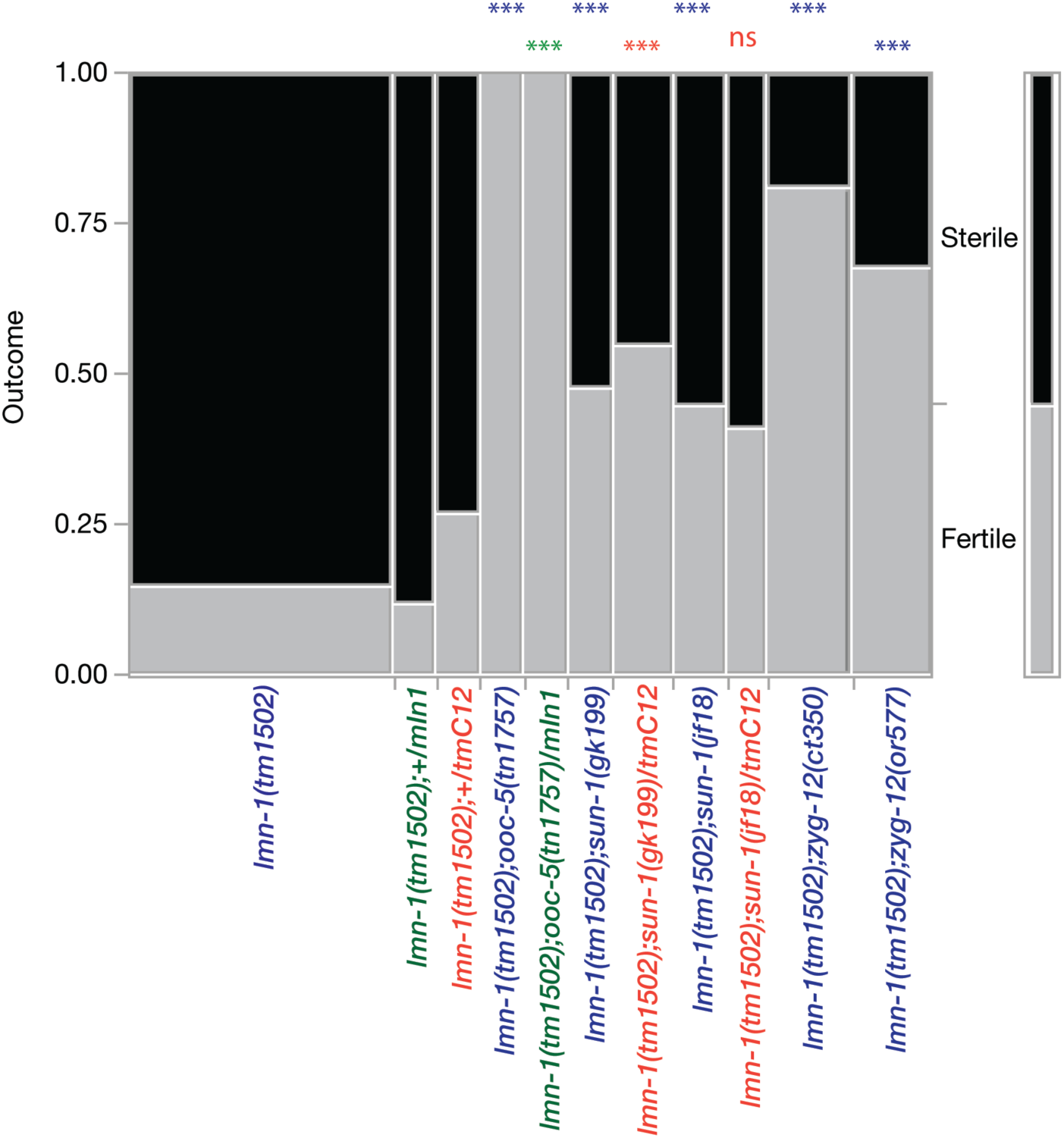
Mutations in *ooc-5* and in the LINC complex components rescue the sterility of *lmn-1* mutants. The observations of the phenotypes depicted in Figure 2, panels A-J were classified into fertile and sterile to perform statistical tests. This table allowed for the contingency analysis of the data. Chi square tests were performed by comparing each strain to its specific control accounting for balancers in the genetic backgrounds analyzed (strains in blue were compared to the corresponding control strain in blue, same with green and orange color coded strains), *** means p<0.001 and ns p=0.1. In addition to the *ooc-5(tn1757)* allele, we observed that the sterility of *lmn-1(–)* was also suppressed by *ooc-5(tn1709)* and *ooc-5(it145)*.

**fig. S3.**
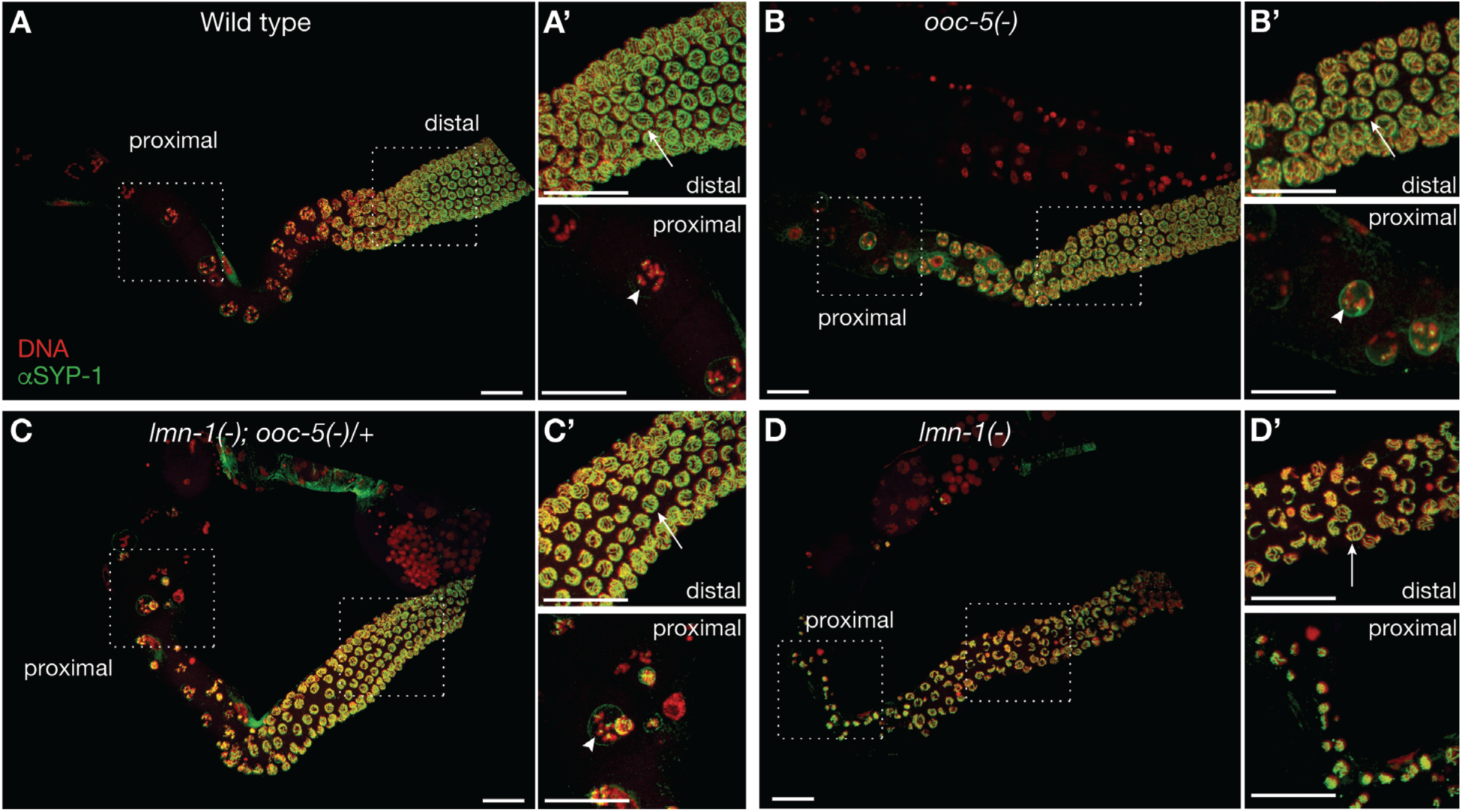
*lmn-1* mutants assemble a synaptonemal complex, but their nuclei collapse after pachytene. Day 1 adult hermaphrodites of the indicated genotypes were dissected, fixed and immunostained for SYP-1(green) and DNA (DAPI, red). (D) The *lmn-1* mutant shown here is an “escaper”, which produced dead embryos (see main text). (A’-D’) Insets are two-fold magnified relative to the main images. Note, synaptonemal complexes are visible in pachytene nuclei (arrows) of *lmn-1(–)* mutants (distal D’), but must nuclei collapse soon after (proximal D’). In contrast, diakinetic chromosomes (arrowheads) are present in *lmn-1(–); ooc-5(–)/+* animals (proximal C’). Bars, 20 µm.

**fig. S4.**
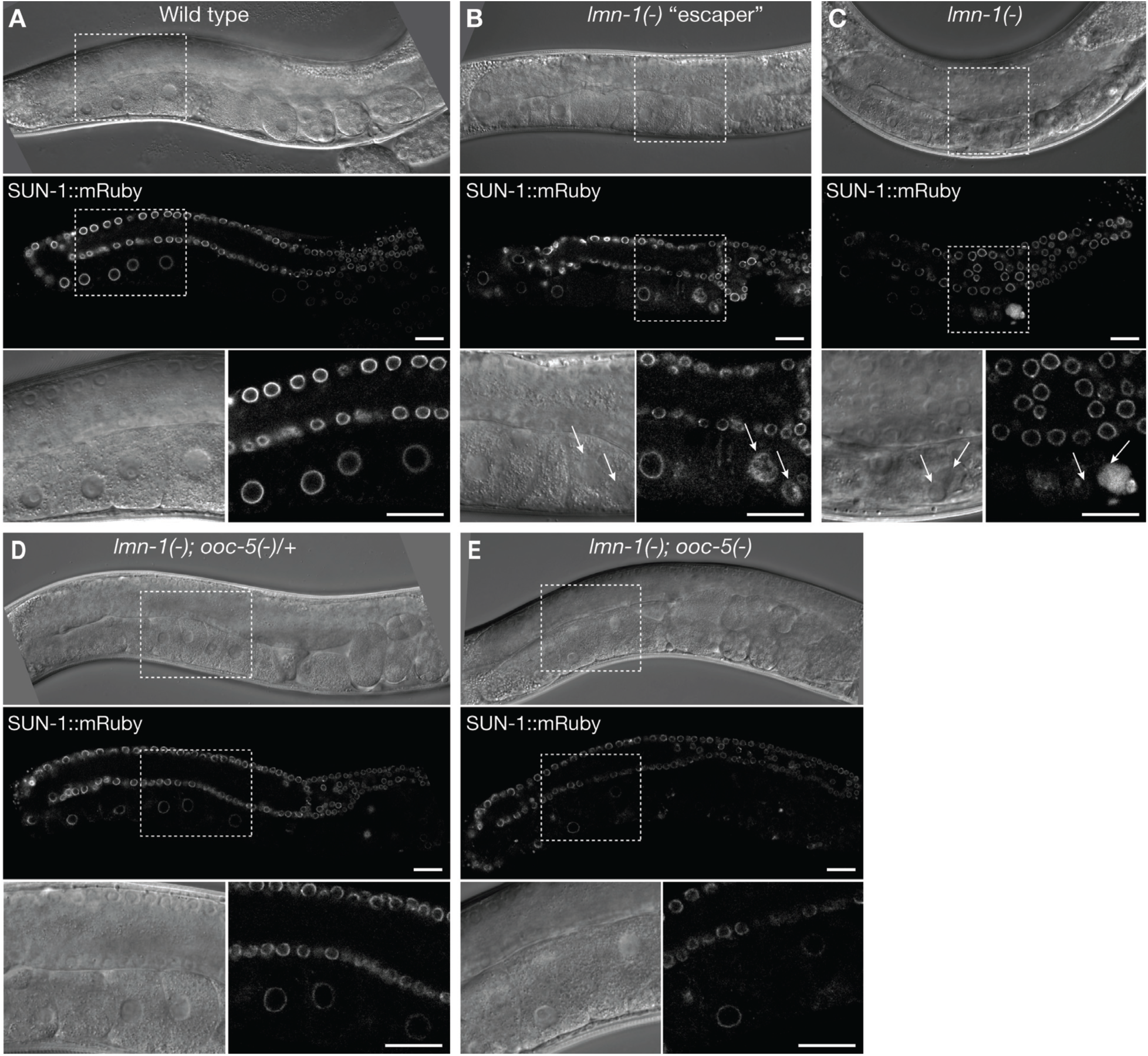
Nuclear damage in *lmn-1* mutants is rescued by reducing *ooc-5* function. (A-E) Confocal and DIC images of the expression of SUN-1::mRuby in the gonad of animals of the indicated genotypes. Animals were grown at 20°C. Insets are two-fold magnified relative to the main images. Arrows in the insets of *lmn-1(–)* mutants (B and C) indicate damaged nuclei with SUN-1::mRuby diffuse localization. Bars, 20 µm.

**fig. S5.**
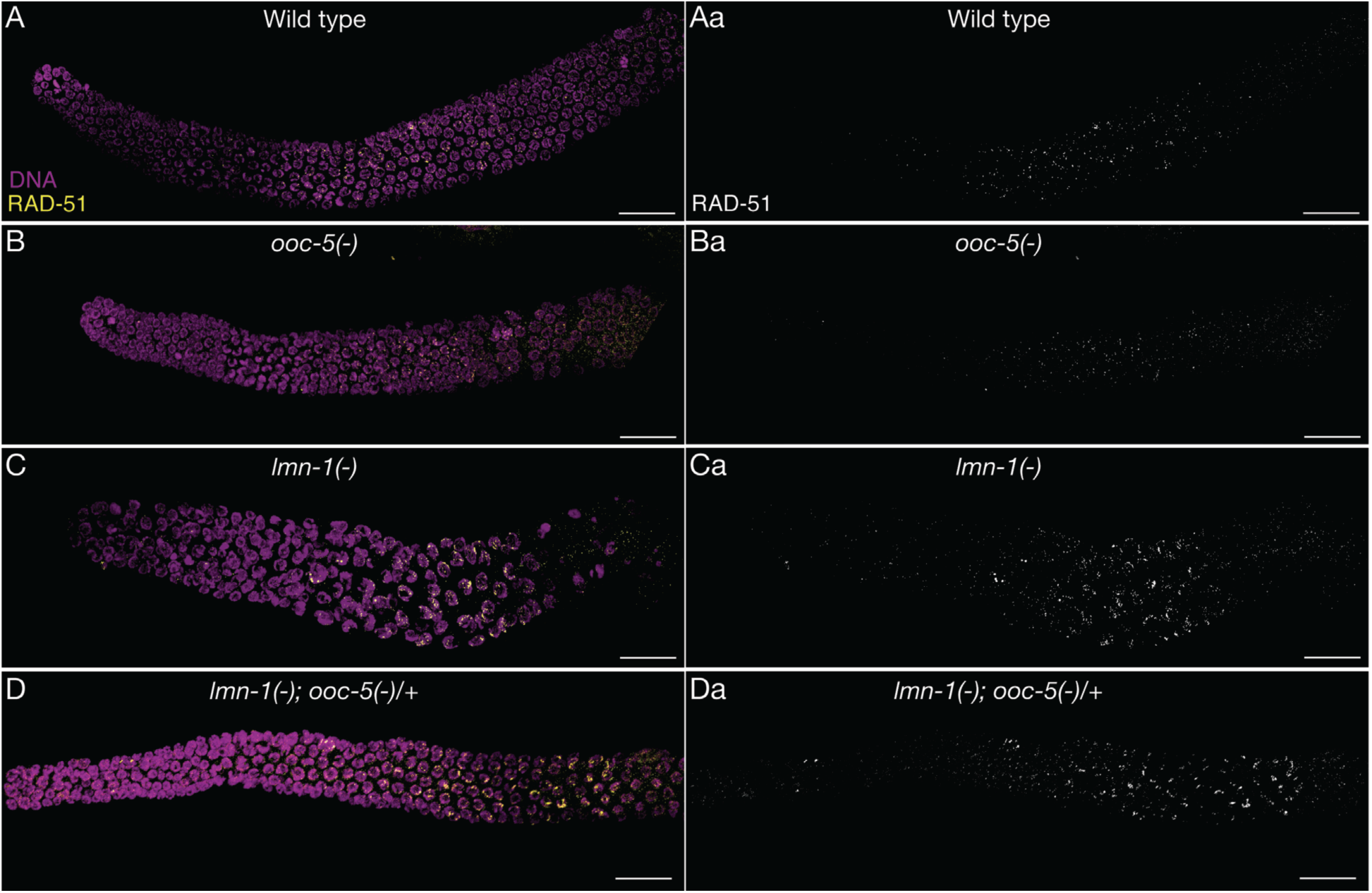
DNA damage in *lmn-1* mutants is not rescued by decreasing *ooc-5* function. (A-D) Day 1 adult hermaphrodites of the indicated genotypes were dissected, fixed and immunostained for RAD-51 (yellow) and DNA (DAPI, magenta). RAD-51 is recruited to sites of DNA double strand breaks to promote successful crossover in normal meiotic cells. (Aa-Da) Color channels were split in A-B to show grayscale images for the RAD-51 staining only. Note, RAD-51 foci increased in size and number in both *lmn-1(–)* (C) and *lmn-1(–);ooc-5(–)/+* (D) mutant strains, suggesting increased DNA damage in these genetic backgrounds. (D) The *lmn-1* mutant shown here is an “escaper,” which produced dead embryos (see main text). Bars, 20 µm.

**fig. S6.**
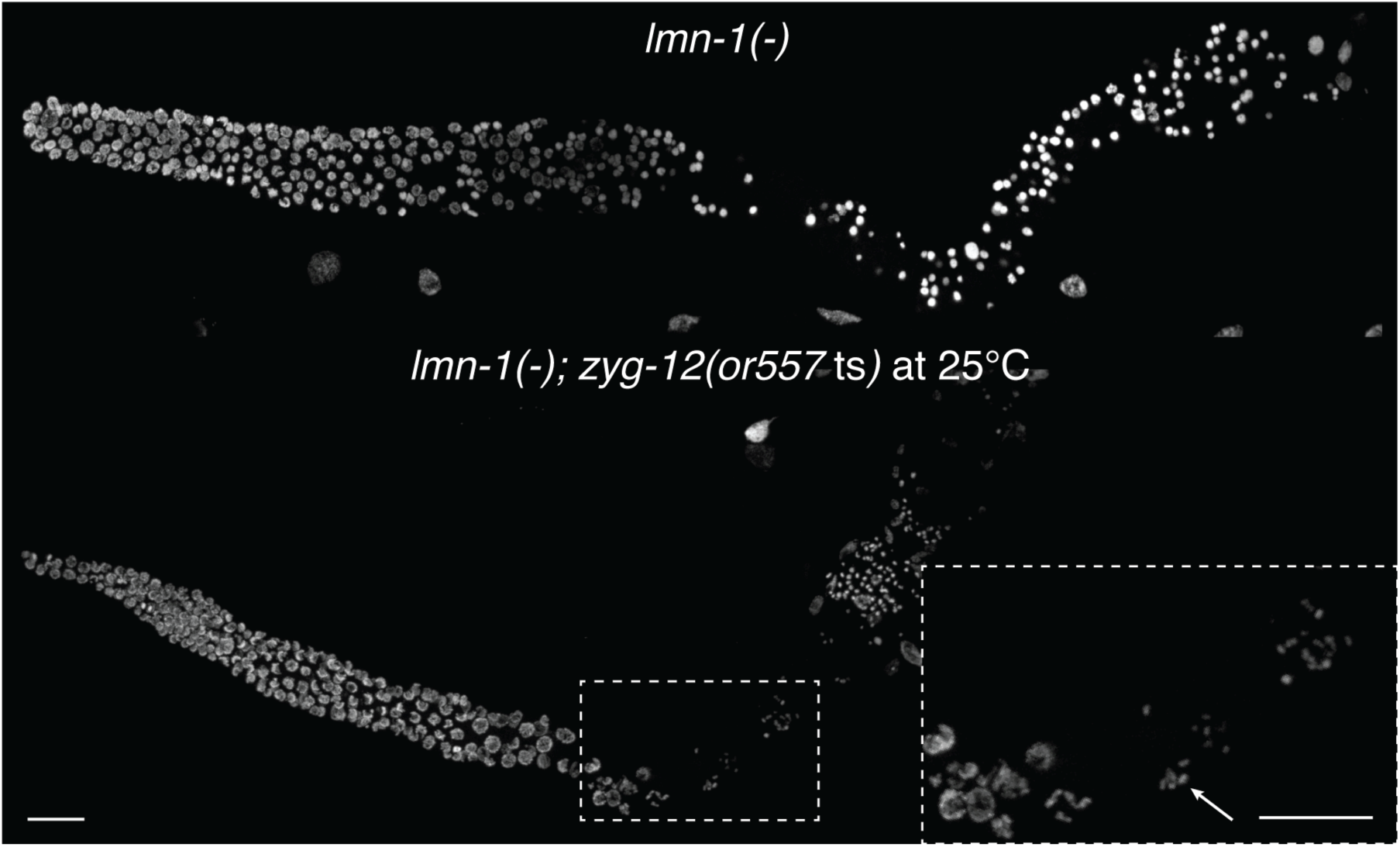
Nuclear damage in *lmn-1* mutants is ameliorated by decreasing LINC complex function. Animals were grown at 20°C until the L4 stage, then *lmn-1(–); zyg-12(or557*ts) double mutants were upshifted to 25°C. *lmn-1(–)* single mutants were kept and analyzed at 20°C because they become sickly when grown at 25°C (unpublished results). Day 1 adult hermaphrodites were dissected, fixed and stained for DNA with DAPI. Note, diakinesis figures are observed in *lmn-1(–); zyg-12(or557*ts) double mutants (arrow) but not *lmn-1(–)* single mutants. Bars, 20 µm.

**fig. S7.**
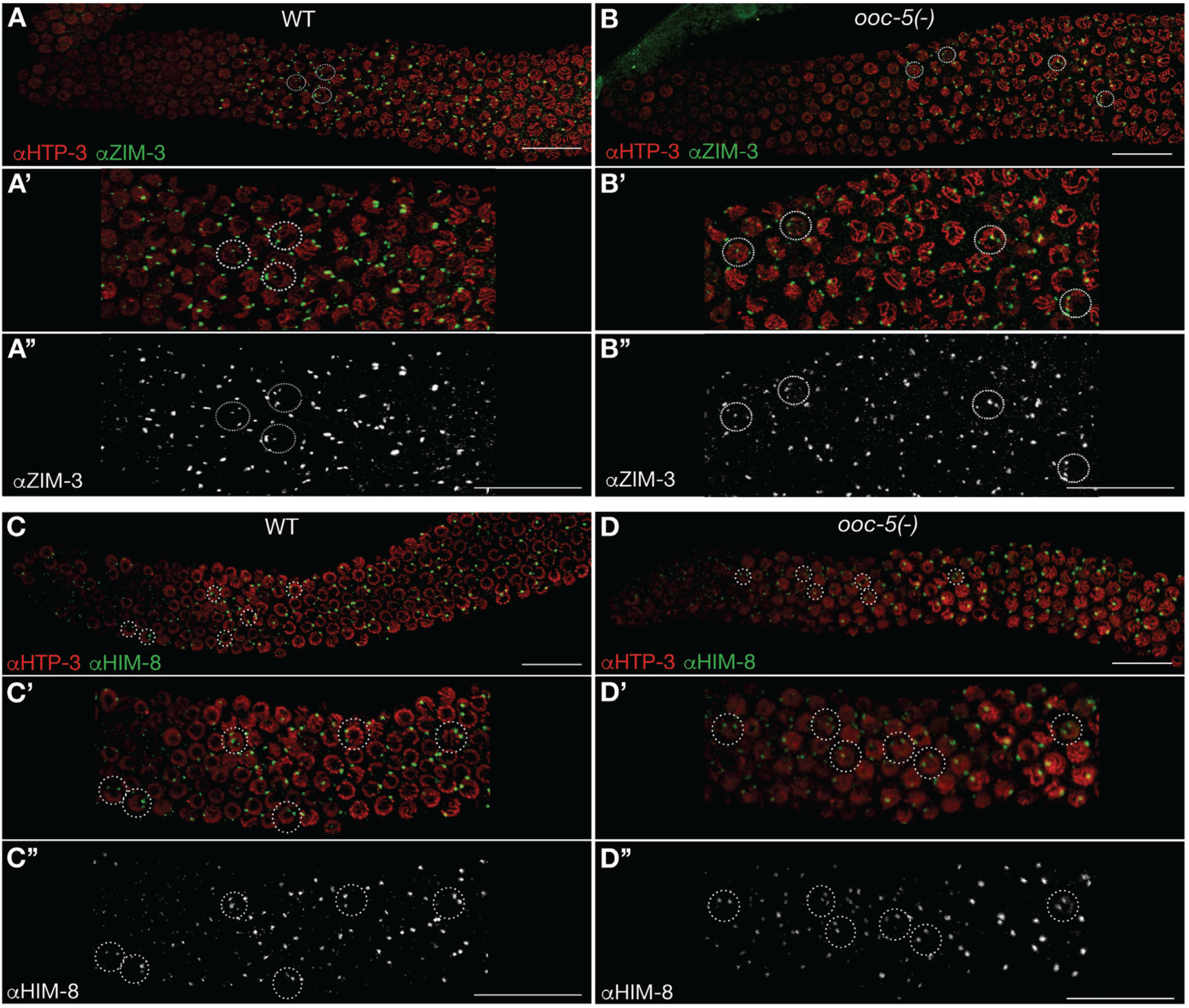
Autosome pairing is delayed in *ooc-5* mutants. Day 1 adult hermaphrodites of the indicated genotypes were dissected, fixed and stained for the axial element protein HTP-3, and either ZIM-3 (A and B) or HIM-8 (C and D) to determine the position of unpaired chromosomes I and IN, and X chromosomes, respectively. Nuclei with unpaired meiotic chromosomes are shown inside the dotted circles. (A’-D’ and A’’-D’’) Insets are two-fold magnified relative to the main images. Bars, 20 µm.

**table S1.**
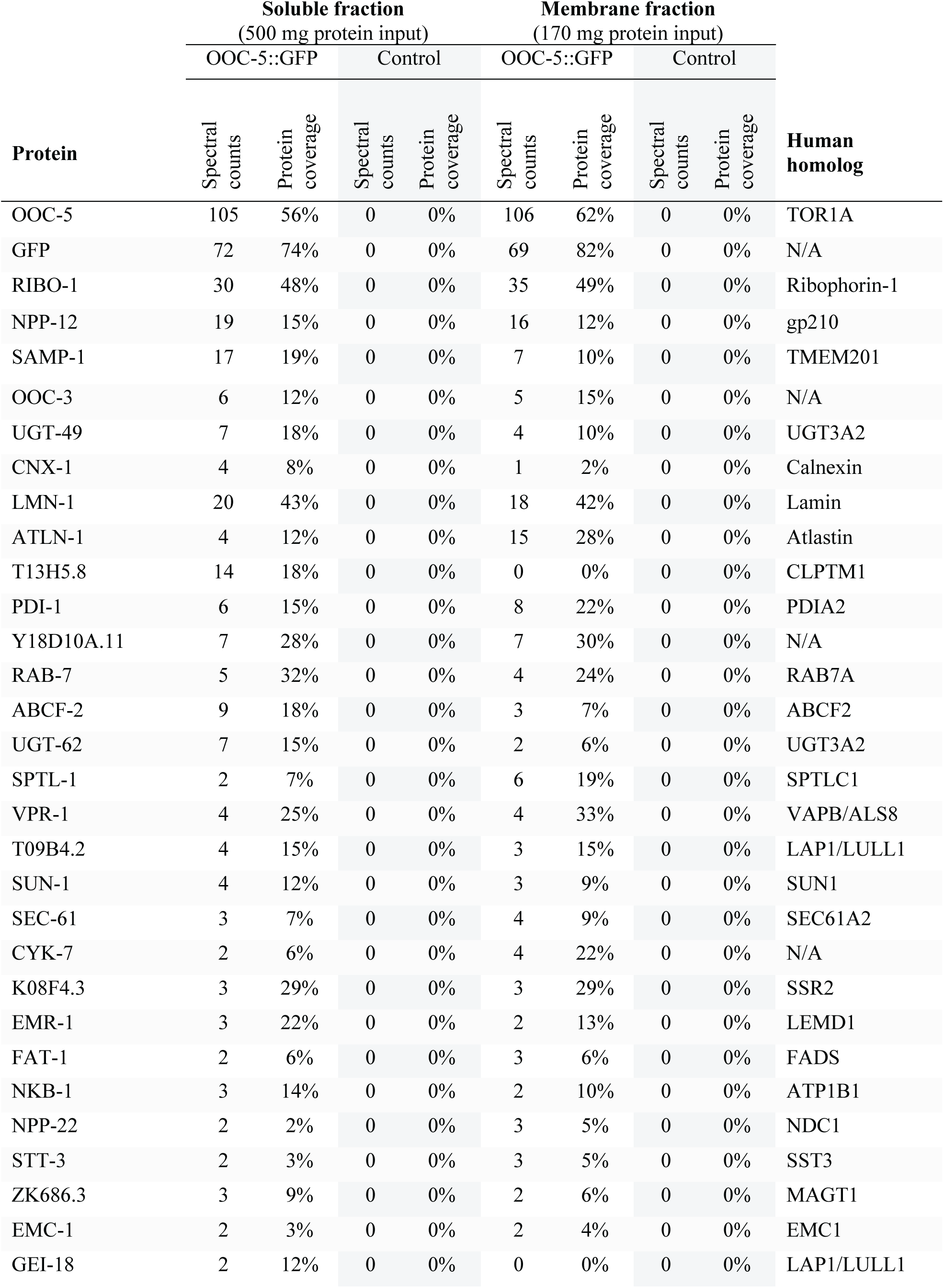
Proteins that co-purify with OOC-5.

**table S2.** Brood sizes. Hermaphrodites were incubated at the indicated temperature (T) throughout their development. Then they were individually selected at the mid-L4 stage and transferred to new plates every 24 h over the course of 3-4 days. Plates were scored for dead embryos, surviving progeny, and males. Embryos not hatching within 24 h after being laid were considered dead.

| Strain | T [°C] | Total eggs laid | Embryonic lethality [%] | Male incidence [%] |
| --- | --- | --- | --- | --- |
| N2 | 15 | 294.9 ± 36.3 | 0.2 ± 0.2 | 0 ± 0 |
|  |  | n=10 | n=2949 |  |
|  | 20 | 290.3 ± 28.6 | 0.7 ± 0.6 | 0.1 ± 0.17 |
|  |  | n=19 | n=5515 |  |
|  | 25 | 288.4 ± 33.7 | 0.29 ± 0.38 | 0.11 ± 0.22 |
|  |  | n=9 | n=2596 |  |
| *ooc-5(tn1757)* | 15 | 81.9 ± 23.4 | 99.2 ± 1.1 | ND |
|  |  | n=30 | n=2457 |  |
|  | 20 | 82.6 ± 20.8 | 98.8 ± 1.8 | ND |
|  |  | n=35 | n=2892 |  |
| *ooc-5(tn1758)*<br>OOC-5::GFP | 15 | 95.7 ± 61.9 | 19.5 ± 14.4 | 0 ± 0 |
|  |  | n=31 | n=2967 |  |
|  | 20 | 64.1 ± 55.4 | 50.6 ± 29.8 | 0 ± 0 |
|  |  | n=29 | n=1860 |  |
|  | 25 | 57.8 ± 39.3 | 80.6 ± 33.3 | 0 ± 0 |
|  |  | n=30 | n=1733 |  |

**table S3.**
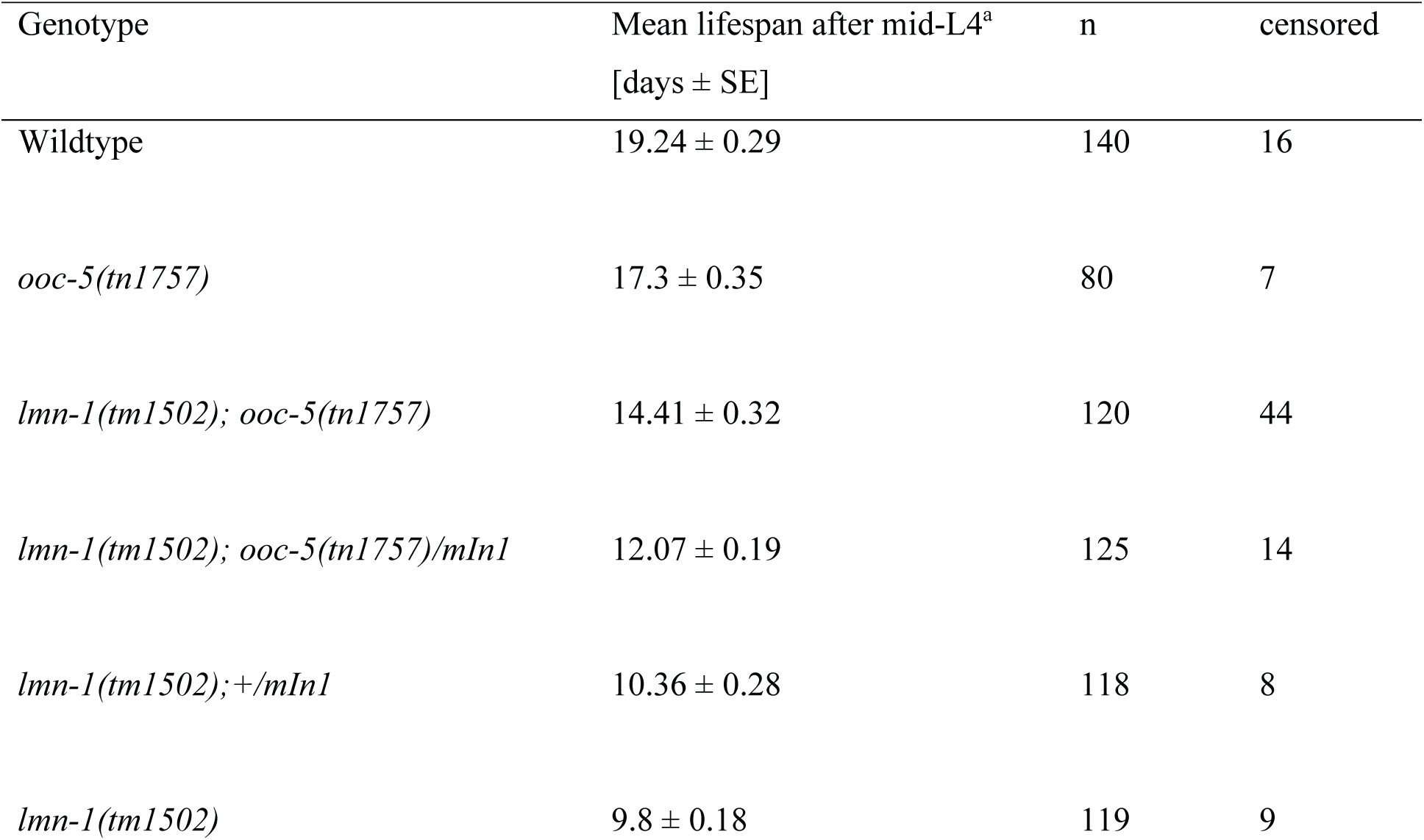
Loss of *ooc-5* extends the lifespan of *lmn-1* mutants. The Wilcoxon tests of individual pairs comparisons for all the genotypes had a p<0.001, except the *lmn-1(tm1502) vs lmn-1(tm1502); +/mIn1* pair; which had a p=0.119. ^a^ For Kaplan-Meier survival curves see Fig. 1E.

**table S4.**
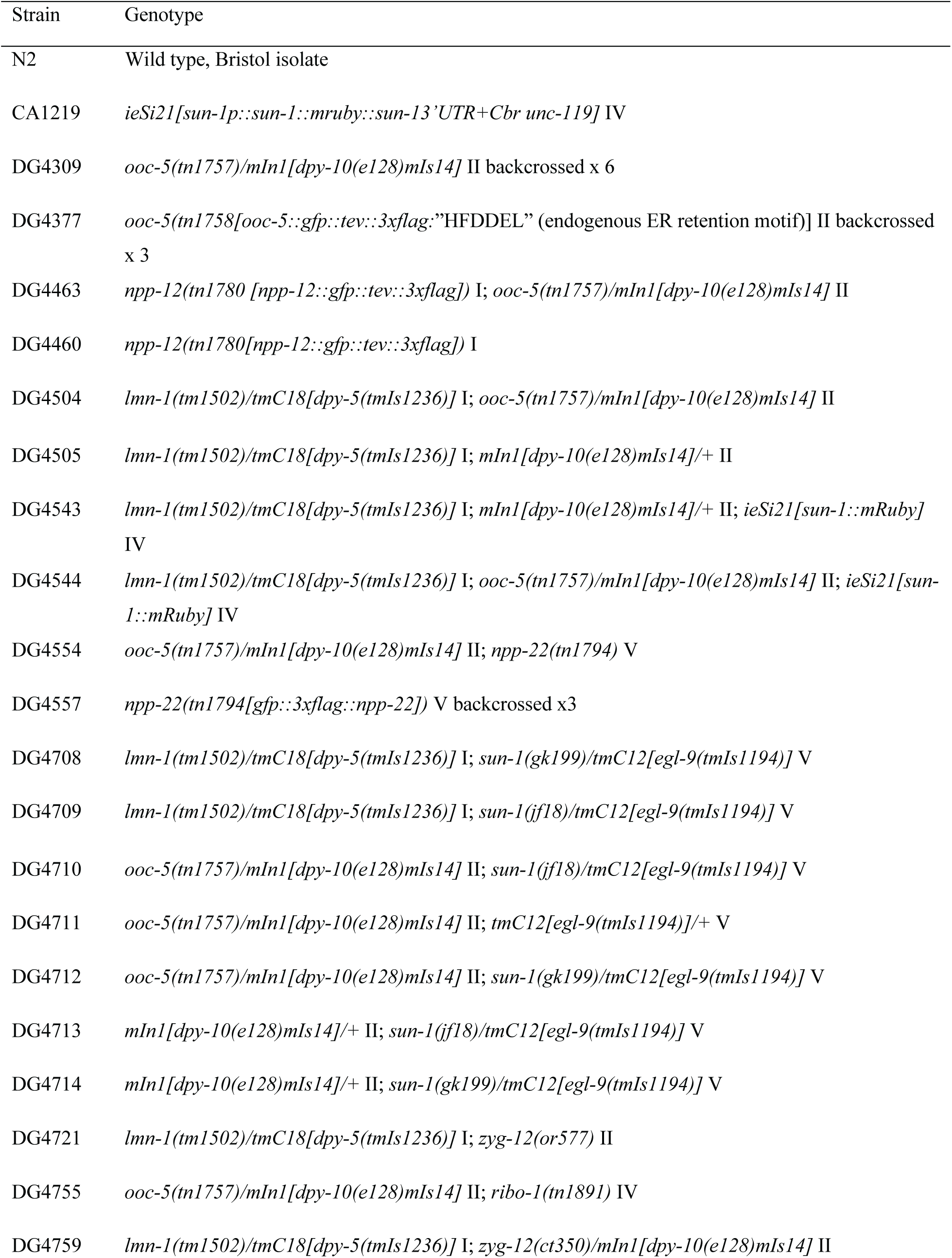
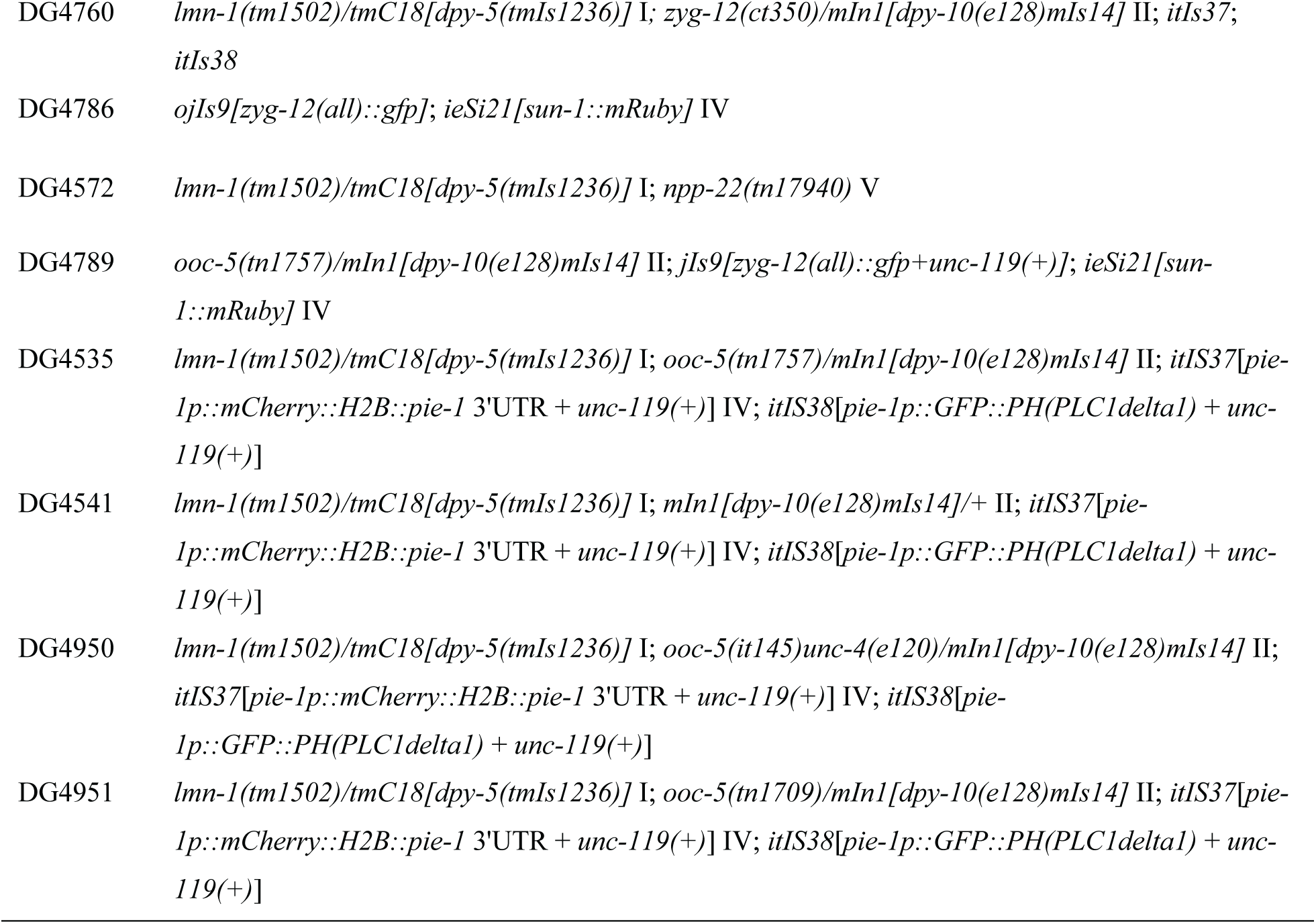
Strains used in this study.

**table S5. Oligonucleotides used in this study.**

**movie S1. Movement of SUN-1 foci in early prophase of wild-type animals.** 1-day old adult hermaphrodites carrying the SUN-1::mRuby expressing transgene were anesthetized and mounted for observation. Z-series were acquired every 15 s for 6-7 min. Maximum intensity projections were converted to grayscale and their LUTs were inverted.

**movie S2. Movement of SUN-1 foci in early prophase of *ooc-5(–)* homozygous animals.** 1-day old adult hermaphrodites homozygous for the *ooc-5(tn1757)* deletion and carrying the SUN-1::mRuby expressing transgene were anesthetized and mounted for observation. Z-series were acquired every 15 s for 6-7 min. Maximum intensity projections were converted to grayscale and their LUTs were inverted.

**movie S3. Movement of SUN-1 foci in early prophase of *ooc-5(–)/+* heterozygous animals.** 1-day old adult hermaphrodites heterozygous for the *ooc-5(tn1757)* deletion over a balancer chromosome (*mIn1*) and carrying the SUN-1::mRuby expressing transgene were anesthetized and mounted for observation. Z-series were acquired every 15 s for 6-7 min. Maximum intensity projections were converted to grayscale and their LUTs were inverted.

